# Structure classification of the proteins from *Salmonella enterica* pangenome revealed novel potential pathogenicity islands

**DOI:** 10.1101/2024.01.30.577995

**Authors:** Kirill E. Medvedev, Jing Zhang, R. Dustin Schaeffer, Lisa N. Kinch, Qian Cong, Nick V. Grishin

## Abstract

*Salmonella enterica* is a pathogenic bacterium known for causing severe typhoid fever in humans, making it important to study due to its potential health risks and significant impact on public health. This study provides evolutionary classification of proteins from *Salmonella enterica* pangenome. We classified 17,238 domains from 13,147 proteins from 79,758 *Salmonella enterica* strains and studied in detail domains of 272 proteins from fourteen characterized *Salmonella* pathogenicity islands (SPIs). Among SPIs-related proteins, 90 proteins function in the secretion machinery. 41% domains of SPI proteins have no previous sequence annotation. By comparing clinical and environmental isolates, we identified 3682 proteins that are overrepresented in clinical group that we consider as potentially pathogenic. 69 of them overlap with SPI proteins. Among domains of potentially pathogenic proteins only 50% domains were annotated by sequence methods previously. Moreover, 36% (1330 out of 3682) of potentially pathogenic proteins cannot be classified into Evolutionary Classification of Protein Domains database (ECOD). Among classified domains of potentially pathogenic proteins the most populated homology groups include helix-turn-helix (HTH), Immunoglobulin-related, and P-loop domains-related. Functional analysis revealed overrepresentation of these protein in biological processes related to viral entry into host cell, antibiotic biosynthesis, DNA metabolism and conformation change, and underrepresentation in translational processes. Analysis of the potentially pathogenic proteins indicates that they form 119 clusters (islands) within the *Salmonella* genome, suggesting their potential contribution to the bacterium’s virulence. Overall, our analysis revealed that identified potentially pathogenic proteins are poorly studied. ECOD hierarchy of classified *Salmonella enterica* domains is available online: http://prodata.swmed.edu/ecod/index_salm.php

**Author Summary:** *Salmonella enterica* is a dangerous bacterium known for causing severe typhoid fever in humans, posing significant health risks. Our study focuses on understanding the proteins of this bacterium’s genetic composition, unraveling their evolutionary classification. Analyzing a vast collection of strains, we identified and classified over 17,000 protein domains, with special focus on 272 proteins within *Salmonella* pathogenicity islands (SPIs). By comparing strains from clinical and environmental sources, we pinpointed 3,682 proteins overrepresented in clinical samples, signifying potential pathogenicity. Surprisingly, half of these proteins’ domains were not identified previously using sequence-based approaches. Our analysis identified these proteins forming 119 clusters within the *Salmonella* genome, suggesting their involvement in its virulence. Our study underscores the insufficient understanding of these potentially pathogenic proteins, highlighting the need for further investigation into their roles and implications in *Salmonella*-related illnesses.

## Introduction

Salmonellae are gram-negative bacteria that are members of the family Enterobacteriaceae. According to the most recent nomenclatural system, there are currently two recognized species: *Salmonella enterica* and *S. bongori* [1]. *S. enterica* comprises six subspecies and accounts for roughly 2000 of the 2600 serovars in the genus [2]. *Salmonella* species can be categorized into typhoid and non-typhoid based on their capacity to induce particular pathologies. Non-typhoid serovars (Typhimurium, Enteritidis, etc.) can cause gastroenteritis and infect humans and other animals [3]. These serovars are primarily spread through animal-derived foods like beef, pork, poultry, and raw eggs that may be contaminated [4] and usually cause mild symptoms that do not require antibiotic treatment [3]. Typhoid serovars (Typhi, Sendai, Paratyphi) have a limited host range, being highly adapted to humans and having only higher primates and humans as their hosts [4]. These serovars lead to the development of typhoid fever and are associated with various symptoms, including high fever, diarrhea, vomiting, headaches, and, in severe cases, death [3]. It is estimated that there are over 20 million cases of typhoid fever annually, leading to more than 200,000 deaths [5]. These serovars can be transmitted through various means, including water, milk, raw vegetables, seafood, and eggs that have been contaminated. For example, S. Typhi and S. Paratyphi do not have animal reservoirs apart from higher primates, their presence suggests contamination resulting from poor hygiene practices during the preparation and handling of food and water [6], and is still a major health problem, especially in the developing world with poor sanitation [7]. Moreover, it was shown that S. Typhi infection increases the risk of gallbladder cancer development [8].

Bacterial secretion systems are intricate molecular machinery found in various bacteria that facilitate the transport of proteins, toxins, and other molecules across the bacterial cell envelope. These systems are crucial for the survival, virulence, and adaptation of bacteria in various environments. There are multiple types of bacterial secretion systems, and they can be broadly categorized into several classes, such as T1SS, T2SS, T3SS, etc. [9]. These systems play a crucial role in bacterial interactions with their environment, including interactions with host cells and other bacteria. Some bacterial secretion systems are associated with virulence and contribute to the bacterial pathogenicity. The proteins responsible for the functioning of bacterial secretion systems are typically found within pathogenicity islands, along with other proteins that enhance the virulence of the bacteria. Pathogenicity islands are a distinct category of genomic islands acquired by microorganisms through horizontal gene transfer. These islands are substantial genomic regions, ranging from 10 to 200 kilobases in size, that become integrated into the genome of pathogenic bacteria and are absent in non-pathogenic bacteria of the same or closely related species [10]. As of today, there are 17 known Salmonella pathogenicity islands, some of which are poorly studied [11]. Understanding secretion systems and the function of pathogenicity-related proteins, as well as the identification of other proteins that contribute to bacterial virulence is essential to develop targeted strategies for combating bacterial infections and diseases.

Recent advancements in protein structure prediction have demonstrated that deep learning approaches like AlphaFold can attain atomic-resolution outcomes, even for such rapidly evolving proteins as bacterial proteins [12]. The structure generated by these structure prediction methods can offer insights into distant homologs that may have lost their sequence signals of homology, thereby providing clues to potential protein functions. The Evolutionary Classification Of protein Domains (ECOD) database [13, 14] provides a valuable resource for analyzing predicted protein structures by placing them in an evolutionary context with experimentally determined protein structures from the Protein Data Bank. ECOD stands as a hierarchical evolutionary classification system that distinguishes itself from other structure-based domain classifications by primarily grouping domains based on homology rather than topology. A significant characteristic of ECOD is its focus on distant homology, culminating in a comprehensive catalog of evolutionary relationships among the categorized domains.

Here we provide an evolutionary classification of protein domains from *Salmonella enterica* pangenome. Using Domain Parser for AlphaFold Models (DPAM) we identified 25,233 domains from 13,147 proteins (protein clusters’ representatives). We evaluated domains 272 proteins from fourteen SPIs and studied their ECOD group distribution based on their relation to the secretion machinery. Moreover, we identified 3682 potentially pathogenic proteins using a comparison of clinical and environmental isolates. The analysis of these proteins indicates that they form 119 clusters or islands within the *Salmonella* genome, suggesting their potential contribution to the bacterium’s virulence. Functional analysis revealed that these proteins are significantly overrepresented in biological processes related to host cell invasion.

## Results and Discussion

### Evolutionary classification of *Salmonella enterica* AlphaFold models

Domains parsing and classification AlphaFold (AF) models of proteins from *Salmonella enterica* pangenome into Evolutionary Classification Of protein Domains (ECOD) [13, 14] hierarchical groups were conducted using Domain Parser for AlphaFold Models (DPAM) [15] recently developed by our group. Overall, DPAM identified 25,233 domains from 13,147 proteins, which are representatives of protein clusters (see Materials and Methods). 17,238 (68%) domains have been assigned to the existing groups in the ECOD hierarchy. The unassigned set consists of 7,995 domains with a low DPAM probability (< 0.85). These domains can either be categorized within the current ECOD hierarchy through manual assessment or may represent new folds that have not been documented in ECOD. ECOD hierarchy of classified *Salmonella enterica* domains is available as a public database, which also provides a downloadable format of the data (http://prodata.swmed.edu/ecod/index_salm.php).

Figure 1 illustrates the distribution of architecture (A-groups) and homology (H-groups) ECOD levels for assigned domains from *Salmonella enterica* pangenome. The most prevalent architecture is a/b three-layered sandwiches, constituting 20% of *Salmonella* domains. This percentage is higher than the fraction (18%) found among classified ECOD domains from experimentally determined structures in the Protein Data Bank (Supplementary Table S1). This A-group includes domains from the proteins that contain Rossmann-like motifs whose function is mostly related to metabolism [16, 17]. The most populated H-group (P-loop domains-related) within this architecture includes ATPases that are part of bacterial secretion systems and play an important role during the injection of bacterial effector proteins [18]. The second most populated A-group, alpha arrays, is the most overrepresented in comparison to ECOD domains from experimental structures. The *Salmonella* set fraction of this architecture (18.7%) is significantly higher than that of the PDB set (9.1%) (Supplementary Table S1). Helix-turn-helix (HTH) domains comprise the majority of alpha arrays A-group (Fig. 1). In bacteria, proteins containing HTH motifs often serve as transcription factors [19]. The “few secondary structure elements” and “extended segments” A-groups are the most significantly underrepresented. These groups contain domains with small and flexible structures that are difficult to classify, contain a limited secondary structure, and might include signal peptides [20].

**Figure 1.**
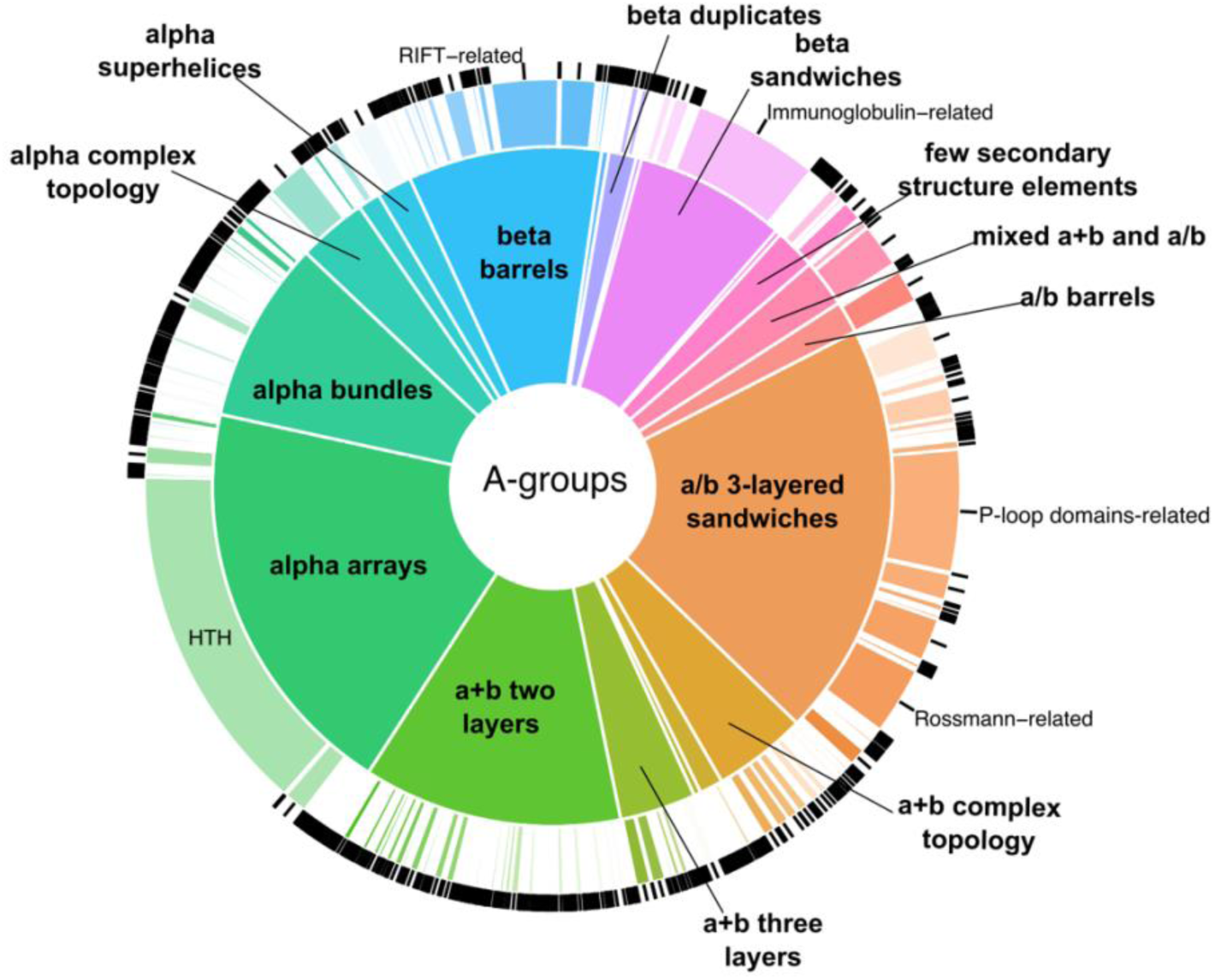
*Salmonella enterica* proteins statistics. Distribution of ECOD A- (inner pie chart) and H-groups (outer pie chart) for domains of AF models from *Salmonella enterica* pangenome.

### Domain classification of *Salmonella* pathogenicity islands (SPIs) related proteins

Using published studies, we created a list of protein-coding genes that are part of *Salmonella* pathogenicity islands (SPIs). Overall, 272 protein-coding genes from SPI-1 – SPI-14 were identified (Supplementary Table S2). SPIs-related proteins were mapped to the representatives of protein clusters retrieved from *Salmonella enterica* pangenome (see Materials and Methods). 28 out of 272 (10%) SPIs-related proteins have experimentally determined 3D protein structure and additionally, 62% (168 out of 272) have homologous proteins among known 3D structures in RCSB Protein Data Bank [21] based on sequence similarity identified by BLAST [22] (E-value < 0.001) (Supplementary Table S2). Thus, 75 out of 272 (28%) of SPIs-related proteins do not have experimentally determined 3D structures and have no close homologs in the RCSB Protein Data Bank. Using DPAM, 223 out of 272 (82%) SPI-related proteins have been classified into ECOD hierarchy up to topology level (T-group). The major reasons for the unclassified 18% of SPIs-related proteins are the high fraction of disordered regions and protein length exceeding 2000 amino acids.

Figure 2 shows the distribution of architecture (A-groups) and homology (H-groups) levels (Fig. 2A) and significantly overrepresented H-groups among SPIs-related proteins in comparison to the *Salmonella* pangenome (Fig. 2B). 41% domains of SPI proteins have no previous sequence annotation and have been determined using AF structural data. The top 3 most populated ECOD A-groups among SPIs related proteins include alpha arrays, a/b three-layered sandwiches and a+b two layers (Fig. 2A). The largest fraction of alpha arrays constitutes HTH H-group. Domains containing helix-turn-helix motifs (HTH) often play an important role in transcription factors in prokaryotes [19, 23]. The most populated SPIs-related H-groups from the a/b three-layered sandwich architecture are P-loop domains-related, Rossmann-related, and Class I glutamine amidotransferase-like. Domains from these H-groups contain minimal Rossmann-like motif (RLM). As we showed previously Rossmann folds are present in ancient domains that have often undergone significant divergence, resulting in a wide array of functions, the execution of various enzymatic reactions, and might constitute up to 30% of the bacterial proteome [16, 17]. Among the significantly overrepresented SPIs-related H-groups, protein domains associated with the type III secretion system (T3SS) are particularly notable (Ring-building motif I in type III secretion system, Type III secretion system domains, Type III secretory system chaperone, etc.) (Fig. 2B). In *Salmonella* T3SS is encoded by SPI-1 and SPI-2, which are the most conserved among *Salmonella* serovars [24, 25].

**Figure 2.**
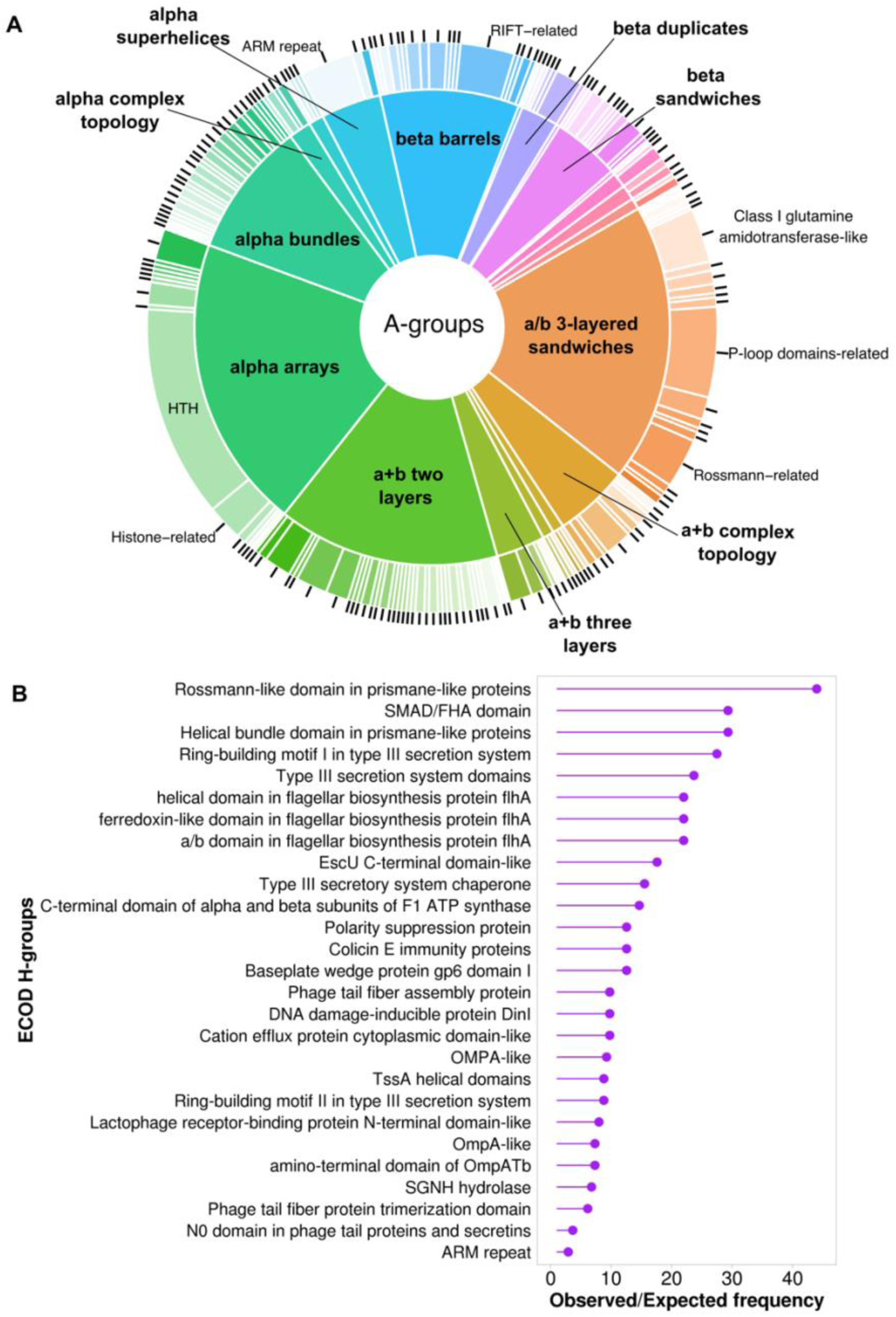
SPIs-related proteins statistics. **(A)** Distribution of ECOD A- (inner pie chart) and H-groups (outer pie chart) for SPIs-related proteins. **(B)** The ratio of observed and expected frequencies of SPIs-related protein domains defines significantly overrepresented (ratio > 1) ECOD H-groups.

Based on published studies we divided SPIs-related proteins into three groups: involved in secretion machinery, not involved in secretion machinery, and unknown function (Supplementary Table S2).

The distribution of ECOD groups for proteins involved in the secretion machinery revealed that A-groups a+b two layers, beta barrels, and alpha bundles have the largest fraction (Fig. 3A). The most populated H-groups from a+b two layers architecture are Ring-building motifs I and II in type III secretion system. Lipoprotein PrgK (UniProt ID: P41786) forms the ring of the injectisome needle complex and contains both these domains [26]. Surface presentation of antigens protein SpaO (UniProt ID: Q56022) adopts two RIFT-related domains from beta barrels architecture. This protein was shown to be the part of sorting platform of type III secretion systems that is responsible for the sequential delivery of the secreted proteins [27, 28]. Intracellular multiplication protein F (IcmF) is an inner membrane protein that forms the core of the type 6 secretion system (T6SS) [29]. This protein is expressed from SPI-6 [30] and contains several structural domains, one of which adopts alpha bundles architecture and is a part of Type VI secretion protein IcmF helical domain ECOD H-group. The distribution of ECOD groups for proteins not involved in the secretion machinery (Fig. 3B) revealed a similarity to the overall distribution of ECOD groups for SPIs-related proteins (Fig. 2A) with a/b three-layered sandwiches, alpha arrays and a+b two layers being the most populated architectures. The A-group a/b three-layered sandwiches also stand out for proteins related to the secretion machinery (Fig. 3A), as this architecture is predominantly represented by a single H-group (P-loop domains-related).

**Figure 3.**
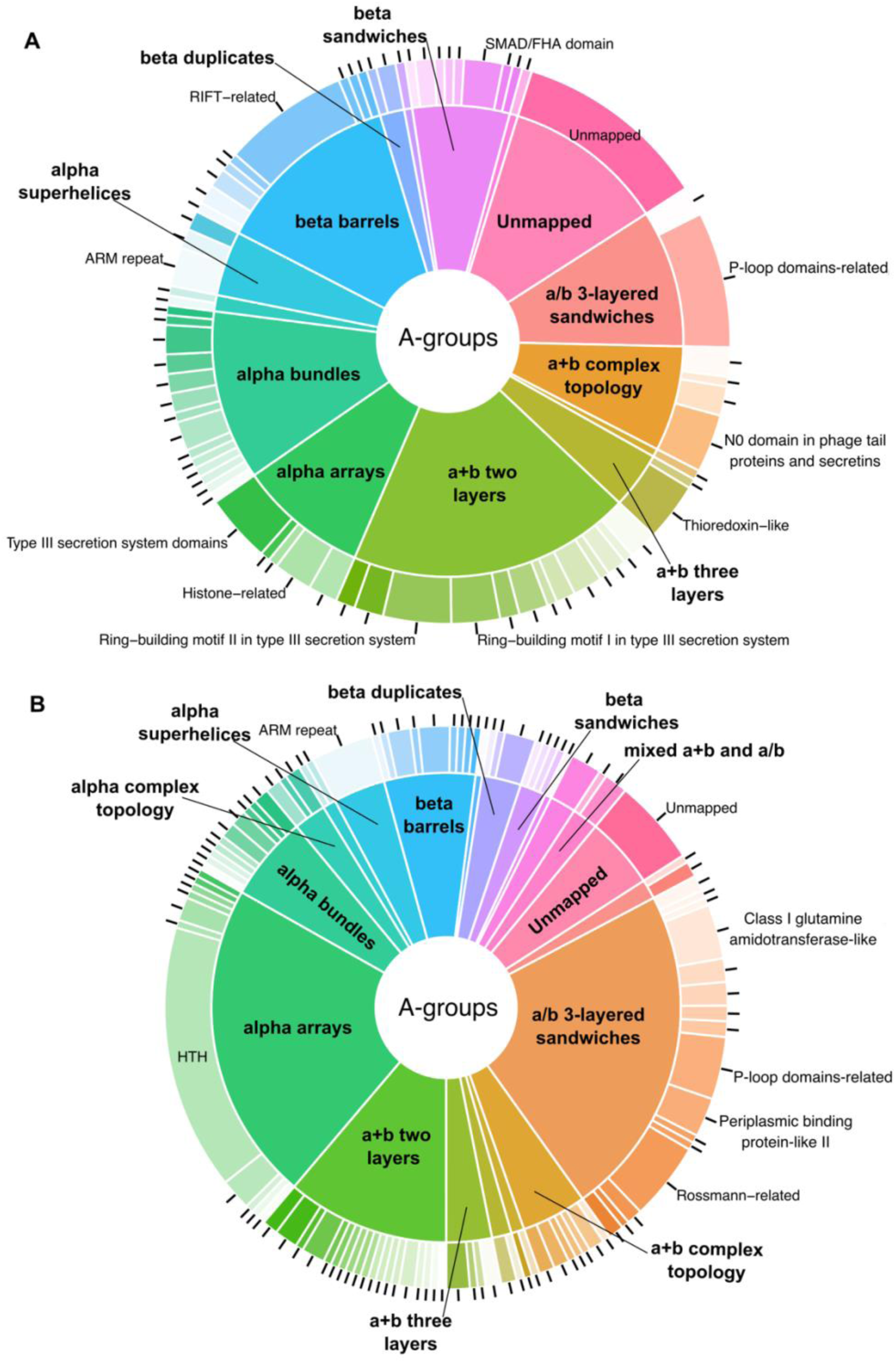
Comparison of SPIs-related proteins involved in secretion machinery functioning vs all rest with known function. **(A)** Distribution of ECOD A- (inner pie chart) and H-groups (outer pie chart) for SPIs-related proteins involved in secretion machinery. **(B)** Distribution of ECOD A- (inner pie chart) and H-groups (outer pie chart) for SPIs-related proteins with known functions not involved in secretion machinery.

SPI proteins that are associated with the secretion machinery and contain P-loop domains serve as ATPases (for example invasion protein InvC, EC: 7.4.2.8, UniProt ID: P0A1B9). These ATPases are integral components of the type III secretion system and are used to inject bacterial effector proteins into eukaryotic host cells [18].

Among all SPIs related proteins only 78% (212 out of 272) contain mapping to the Pfam database [31] (Supplementary Table S2). Our analysis using HHPred [32] showed that some of the 22% unmapped proteins contain domains that have confident hits in the Pfam database. For example, secretion system apparatus protein SsaP (UniProt: P0A239), which is part of the T3SS secretion machinery, showed a similarity with two Pfam families: PF02120 - Flagellar hook-length control protein FliK (Probability: 97.88%, E-value: 2.0E-4) and PF09483 - Type III secretion protein (HpaP) (Probability: 97.84%, E-value: 9.5E-5). The inner membrane protein fidL (UniProt: H9L432) showed similarity to PF15941 - FidL-like putative membrane protein (Probability: 99.78%, E-value: 1.8E-18) and pilus assembly protein pilP (UniProt: Q9RHF2) showed similarity to PF17456 - Toxin-coregulated pilus protein S (Probability: 99.79%, E-value: 3.5E-18). Among all SPIs-related proteins, 36% (99 out of 272) don’t have Gene Ontology assignments according to UniProt KB (Supplementary Table S2) and their function is often unclear. For example, the Inner membrane protein (UniProt: H9L4B1, gene: orf319), which is located within SPI-2 [24], has no GO assignment and its function is vague. The N-terminal domain of H9L4B1 adopts a Flavodoxin-like structure that belongs to the Class I glutamine amidotransferase-like homology group (Fig. 4A). Structure searches against Protein Data Bank using FoldSeek [33] revealed [4Fe- 4S]-dependent thiouracil desulfidase TudS from *E. coli* (PDB ID: 6ZW9) as the most similar (Fig. 3B). This enzyme catalyzes desulfuration of thiouracil to uracil and harbors a [4Fe-4S] cluster bound by three cysteines colored in magenta (Fig. 4B) [34]. H9L4B1 contains three cysteines at the same positions as TudS and one additional cysteine at the C-terminal part of the third beta strand (Fig. 4A). This suggests that H9L4B1 is also capable of binding iron-sulfur clusters and might have potential enzymatic activity. Moreover, both these domains are members of the Pfam family PF04463 (2-thiouracil desulfurase). Another example is the ParB family protein (UniProt: Q8Z1M1) whose function is also unclear. ParB structure includes three domains, two of which have been classified by DPAM. The N-terminal domain, shown in light blue (Fig. 4C), adopts a+b complex topology and is affiliated with the ParB/Sulfiredoxin homology group. The middle domains, shown in light orange, adopt an alpha arrays architecture and belong to the Histone-related H-group. For the C-terminal domain, depicted in red (Fig. 4C), DPAM did not identify a confident hit in ECOD. FoldSeek search against AFDB using the C-terminal domain of Q8Z1M1 showed many bacterial proteins with this domain. For example, Glutamate-1-semialdehyde 2,1-aminomutase from *Nocardia brasiliensis* (UniProt: K0EV93) (Fig. 4D). This protein contains two domains: C-terminal which is similar to the C-terminal domain of ParB, and N-terminal, which belongs to Pfam family Aminotransferase class-III (PF00202). Several known experimental protein structures belong to PF00202, one of them is dialkylglycine decarboxylase from *Burkholderia cepacian* (PDB: 1D7R) (Fig. 4E). The C-terminal domain of this protein belongs to C-terminal domain in some PLP-dependent transferases ECOD H-group. Therefore, C-terminal domains of ParB, glutamate-1-semialdehyde 2,1-aminomutase, and dialkyl glycine decarboxylase are homologous, however, our classification revealed topological differences – decarboxylase contains an additional beta hairpin shown in magenta (Fig. 4F-H).

**Figure 4.**
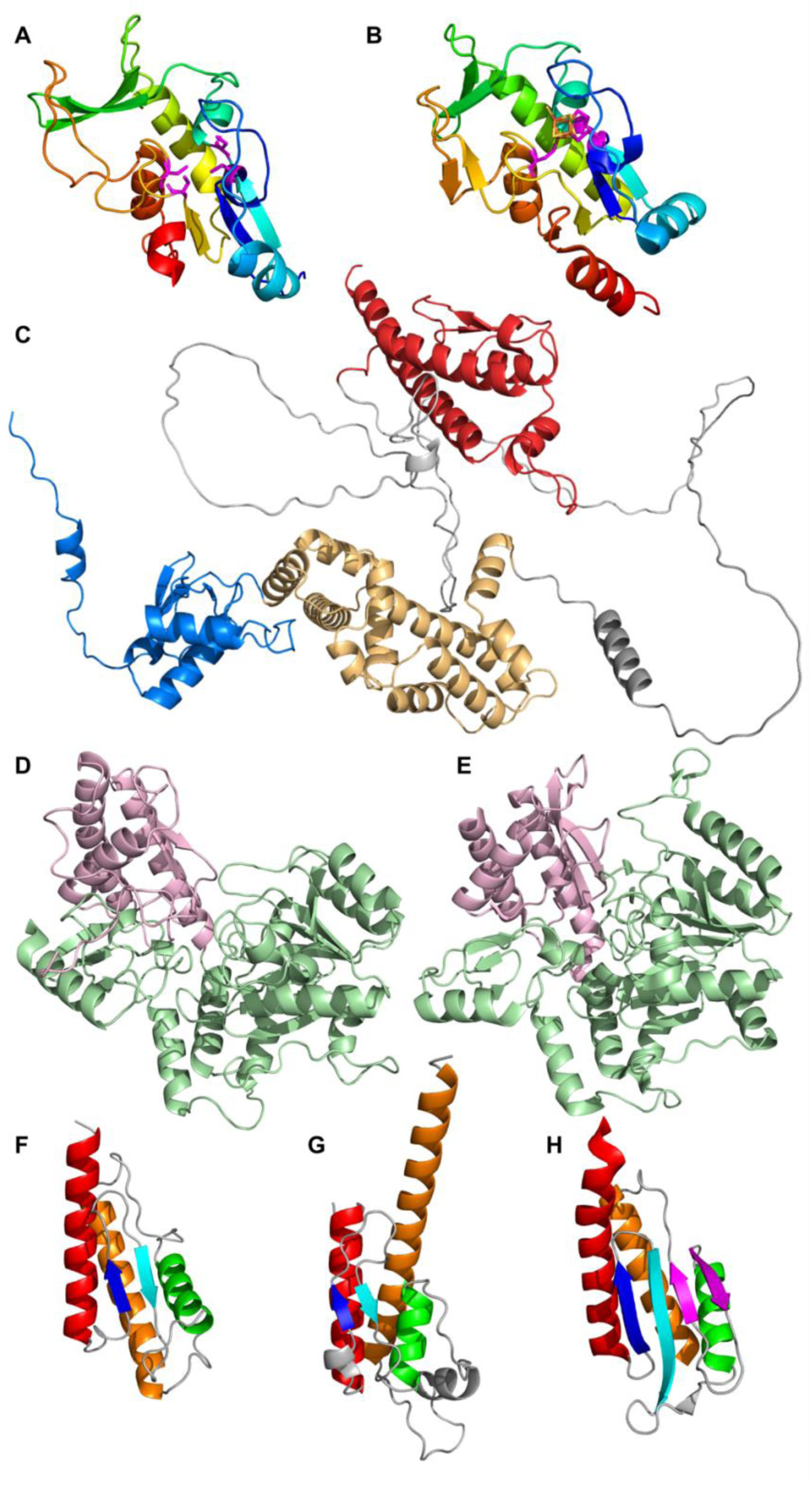
Representative domain structures of SPIs-related proteins. **(A)** N-terminal domain of Inner membrane protein (UniProt: H9L4B1, gene: orf319). Cysteines are colored magenta. **(B)** [4Fe-4S]-dependent thiouracil desulfidase TudS from *E. coli* (PDB: 6ZW9). Cysteines are colored magenta. **(C)** ParB family protein structure (UniProt: Q8Z1M1). **(D)** Glutamate-1-semialdehyde 2,1-aminomutase from *Nocardia brasiliensis* (UniProt: K0EV93). The N-terminal domain is shown in light green, C-terminal in light pink. **(E)** Dialkylglycine decarboxylase from *Burkholderia cepacian* (PDB: 1D7R). The N-terminal domain is shown in light green, C-terminal in light pink. **(F)** C-terminal domain of Glutamate-1-semialdehyde 2,1-aminomutase (UniProt: K0EV93). **(G)** C-terminal domain of ParB family protein (UniProt: Q8Z1M1). **(H)** C-terminal domain of dialkylglycine decarboxylase (PDB: 1D7R).

### Potentially pathogenic proteins revealed similarity in ECOD group distribution with SPIs-related proteins

We defined potentially pathogenic proteins in *Salmonella enterica* pangenome using a comparison of patients and all other types of isolates (food, environment). The least conserved proteins between these two groups were considered potentially pathogenic. Identified proteins were clustered to remove redundancy (see Materials and Methods) which resulted in 3682 clusters of potentially pathogenic protein clusters. Representative proteins from these clusters were used for further analysis. To verify the enrichment of known virulence-related proteins among potentially pathogenic proteins we ran BLAST [22] against the virulence factor database (VFDB) [35] and calculated: 1) observed frequency of virulence-related proteins in the set of identified proteins and 2) expected frequency of virulence-related proteins in *Salmonella enterica* pangenome. The ratio of observed and expected frequencies was 1.282 with a Chi-Square test P-value < 0.00001. It means that virulence-related proteins are significantly overrepresented in the set of identified potentially pathogenic proteins in comparison to the whole pangenome.

We mapped potentially pathogenic proteins to the UniProt database and retrieved Gene Ontology (GO) annotations. Only 45% of these proteins have GO annotations. We calculated over and underrepresentation of biological process categories from the GO Metagenomic subset for potentially pathogenic proteins. Our analysis revealed that these proteins are overrepresented in biological processes related to viral entry into host cell, antibiotic biosynthesis, DNA metabolism and conformation change, and underrepresentation in translational processes (Fig. 5).

**Figure 5.**
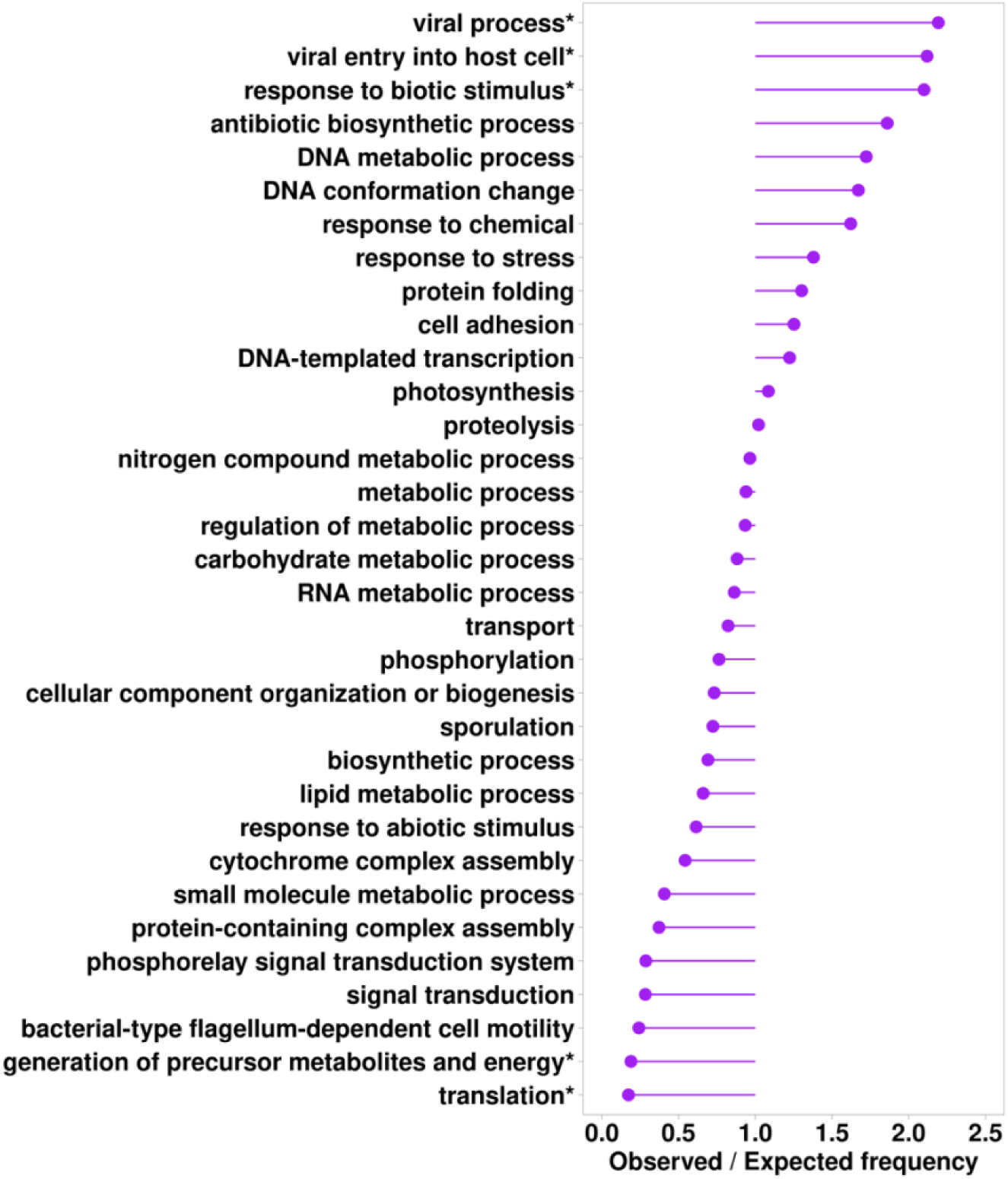
The ratio of observed and expected frequencies of biological processes from the GO Metagenomics subset defines over (ratio>1) and under (ratio<1) represented biological process categories for potentially pathogenic proteins. Asterisks denote significant values according to the chi-square test (P<0.0001).

Distribution of ECOD architecture and homology groups of potentially pathogenic proteins (Fig. 6A) showed significant similarity with one of the SPIs-related proteins (Fig. 2A). Interestingly, 50% of these proteins’ domains have not been annotated previously using sequence-based approaches in the InterPro database [36]. In both cases the dominating A-groups are alpha arrays, a/b three-layered sandwiches, a+b two layers, alpha bundles, and beta barrels. We consider this similarity as further confirmation of the virulence association of identified potentially pathogenic proteins. The most notable difference in the distribution of ECOD groups between SPIs-related and potentially pathogenic proteins is the significantly higher proportion of Immunoglobulin-related (beta sandwiches architecture) and Restriction endonuclease-like (a/b three-layered sandwiches architecture) H-groups in potential pathogenic proteins. These two H-groups are among significantly overrepresented groups in potential pathogenic proteins in comparison to the whole pangenome (Fig. 6B). Restriction endonuclease-like homology group is represented by single ECOD T-group (ECOD ID: 2008.1.1). Restriction endonucleases or restrictases are bacterial and archaeal enzymes that cleave foreign DNA (for example bacteriophages) into fragments as a defense mechanism against invading viruses [37]. Immunoglobulin-related homology group is represented by three T-groups in our dataset. T-group Immunoglobulin/Fibronectin type III (ECOD ID: 11.1.1) includes domains from such *Salmonella* proteins as alpha-2-macroglobulin (PDB: 4U48), which is a plasma membrane protein and protects the bacterial cell from host peptidases [38]. The second T-group Prealbumin-like (ECOD ID: 11.1.4) includes such proteins with experimentally defined structures as lipoprotein TssJ from *Pseudomonas aeruginosa* (PDB: 3ZHN) which is an outer-membrane-associated lipoprotein and is crucial for the function of type VI secretion system [39]. Finally, the third T-group Common fold of diphtheria toxin (ECOD ID: 11.1.5) includes *Salmonella* fimbrial protein SefD with experimentally defined structure (PDB: 3UIY). It was shown that SefD is required for uptake or survival in macrophages after infection [40]. Therefore, the examples discussed above collectively represent groups of proteins that can contribute to *Salmonella* pathogenicity.

**Figure 6.**
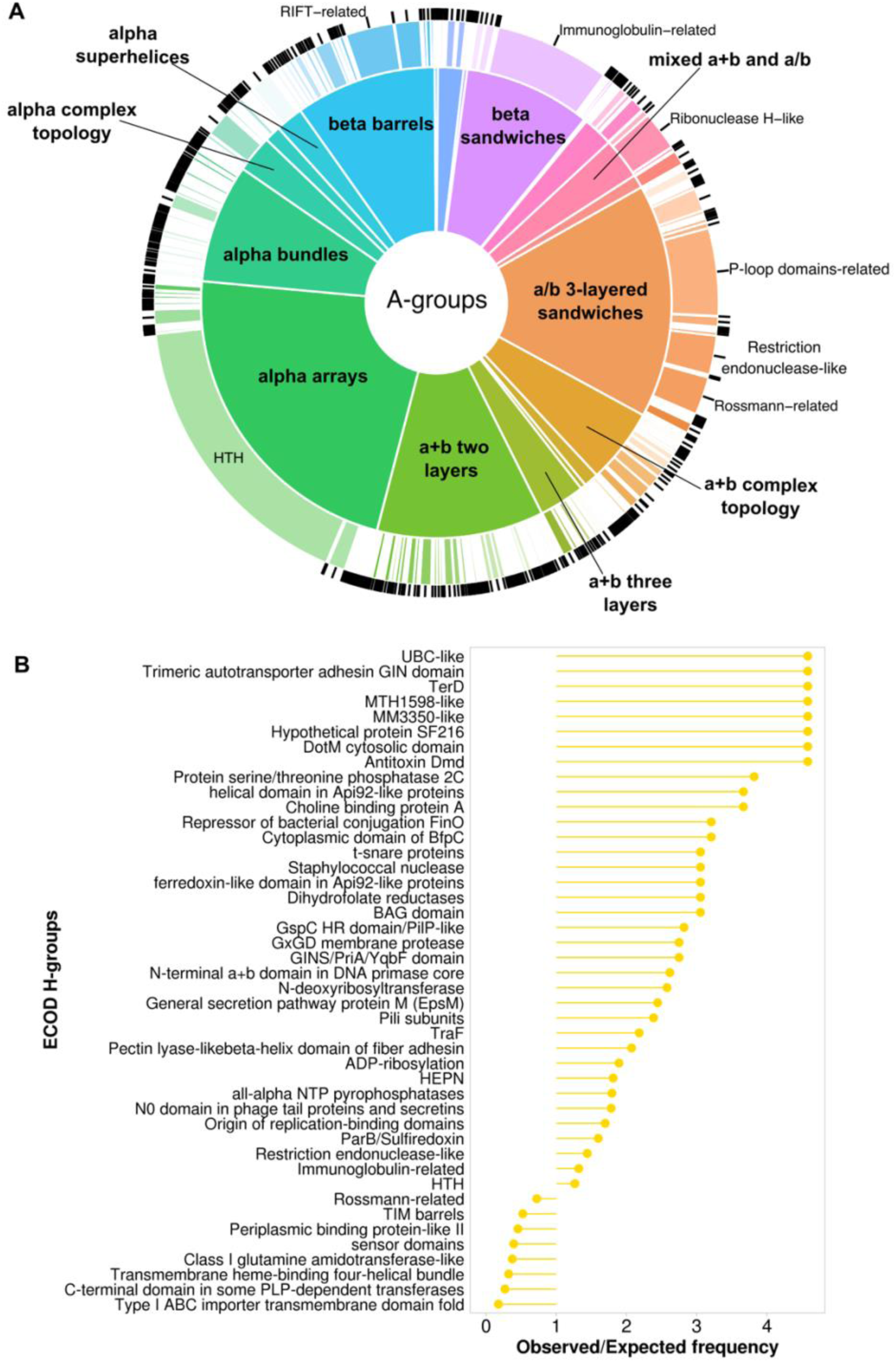
Potentially pathogenic proteins statistics. **(A)** Distribution of ECOD A- (inner pie chart) and H-groups (outer pie chart) for potentially pathogenic proteins. **(B)** The ratio of observed and expected frequencies of potentially pathogenic protein domains defines significantly overrepresented (ratio > 1) and underrepresented (ratio < 1) ECOD H-groups.

For representatives of each cluster from the whole *Salmonella enterica* pangenome we calculated the average pair-wise distance and using Multidimensional Scaling (MDS) we converted the pair-wise distance matrix to one dimension. By sorting representatives by MDS value we obtained the list of protein-coding genes, in which adjacent genes should also be adjacent in the genome. Using chosen criteria (see Materials and Methods) we identified 119 novel potential pathogenicity islands (NPPIs) (Supplementary File 1). One of the NPPIs (NPPI99) revealed significant overrepresentation (ratio of observed and expected frequencies = 6.15; Chi-Square P-value = 3.77E-08) of potentially pathogenic proteins with confident hits within VFDB. Distribution of ECOD groups of potentially pathogenic proteins within NPPI99 revealed the dominance of homology groups related to type 3 secretion system and pilus elements (Fig. 7).

**Figure 7.**
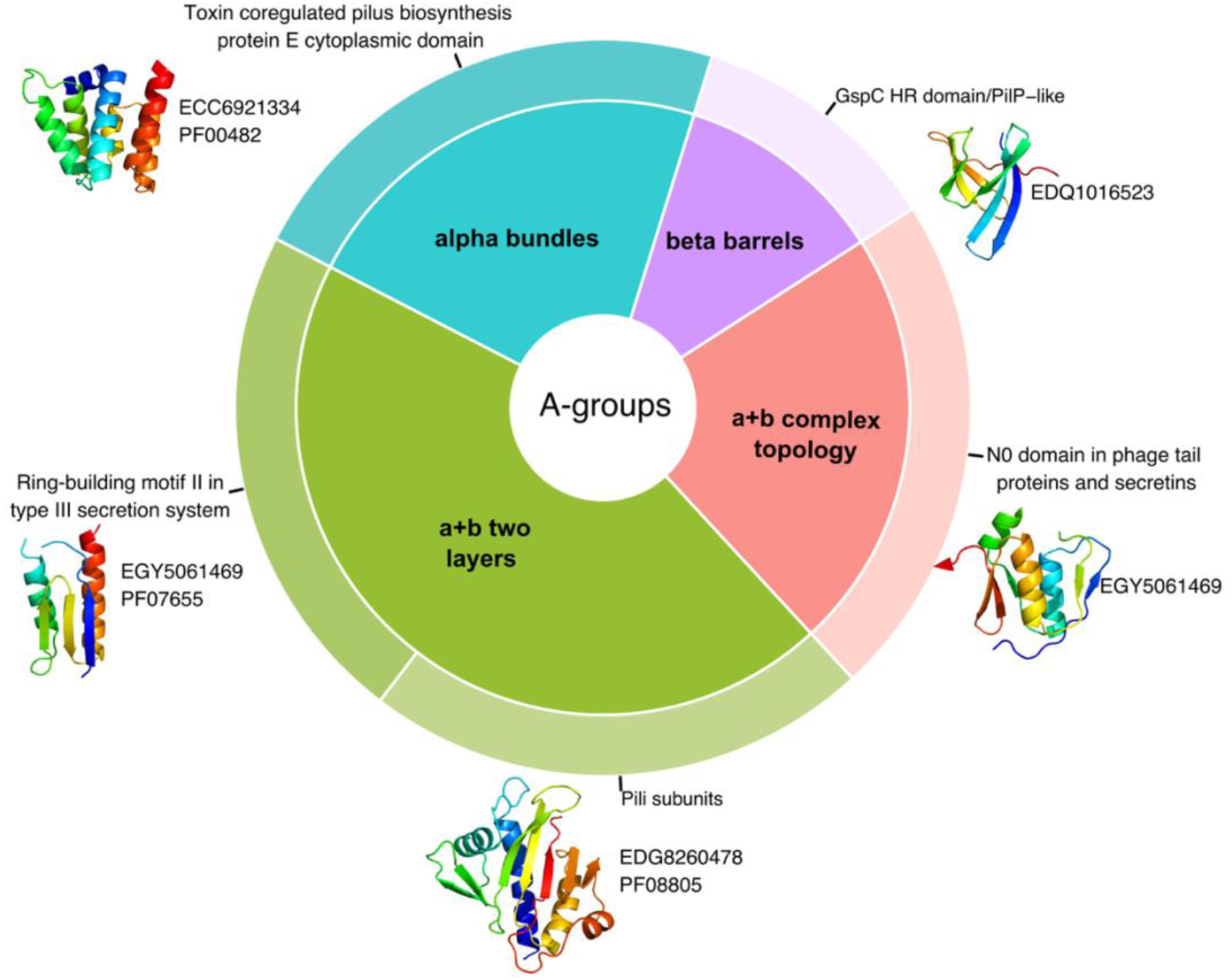
Novel potential pathogenicity island (NPPI99) proteins statistics. Distribution of ECOD A- (inner pie chart) and H-groups (outer pie chart) for potentially pathogenic proteins within NPPI99. NCBI and Pfam IDs are shown near domain structures.

However, the majority of the identified NPPIs do not feature many proteins with confident hits within VFDB and include a significant number of proteins with unknown functions. An example of this is NPPI7, which contains 50% (9 out of 18) of potentially pathogenic proteins with unclear functions (Supplementary Table S3). The distribution of ECOD categories within NPPI7 revealed the prevalence of alpha arrays, beta barrels, and a+b two-layer architectures (Fig. 8A). The most populated A-group alpha arrays are represented by a single H-group – HTH. Domains containing helix-turn-helix motif (HTH) are known to be part of proteins that often function as transcription factors [19]. Indeed, one of the proteins from NPPI7 that consist of one HTH domain is a helix-turn-helix transcriptional regulator (NCBI: ECO0783223.1) (Fig. 8B). Hypothetical protein K5A74_004162 with unknown function (NCBI: EIB4898420.1) adopts two HTH domains (Fig. 8C). The best HHpred hit against Pfam database (Probability: 97.79%, E-value: 0.000026) revealed Domain of unknown function (DUF1870, PF08965) for both domains, which is part of HTH Pfam clan. BLAST run against UniProt KB revealed the set of nearly identical bacterial proteins XRE family transcriptional regulator (Identity: 97.5%, E-value: 1.9e-105). Therefore, the hypothetical protein K5A74_004162 is likely a DNA-binding protein involved in transcription regulation. Hypothetical protein AAK88_22130 (NCBI: ECO0783211.1) adopts a+b two layers topology and is assigned to Glyoxalase/Bleomycin resistance protein/Dihydroxybiphenyl dioxygenase H-group (Fig. 8D). Our analysis showed that this protein belongs to Pfam family DUF5983 (Family of unknown function, PF19419). Its closest homolog identified by DPAM is Toxoflavin Lyase (TflA) from the bacteria *Paenibacillus polymyxa* (PDB: 3PKX). TflA is responsible for the degradation of the toxin toxoflavin [41]. The residues that take part in toxoflavin binding are shown in magenta (Fig. 8E). Hypothetical protein AAK88_22130 contains four residues (two tryptophans, one arginine, and one aspartic acid) that might coordinate a small molecule in a similar position to how TflA coordinates toxoflavin (Fig. 8D-E). Hypothetical protein QD83_004322 (NCBI: EDV2320831.1) adopts beta barrels architecture and is assigned to the SH3 H-group. This protein includes one beta barrels domain and a single alpha helix at the N-terminal end (Fig. 8F). The closest homolog identified by DPAM is bacterial translation initiation factor 5A (IF-5A, PDB: 1BKB) (Fig. 8G). IF-5A plays a role in the initial stage of peptide bond formation during translation [42]. HHpred run against the Pfam database for the hypothetical protein QD83_004322 revealed a significant hit (Probability: 96.95%, E-value: 0.00031) only for N-terminal helix - lysis protein (PF02402). This family includes a signal peptide motif and a lipid attachment site. The beta-barrel domain of QD83_004322 does not have significant hits in the Pfam database. BLAST run against UniProt KB revealed the set of bacterial lipoproteins with 98% sequence identity (E-value: 3.4e-91; UniProt: A0A626QDK6). Thus, the hypothetical protein QD83_004322 is likely a lipoprotein with a lipid attachment site as defined by the Pfam family.

**Figure 8.**
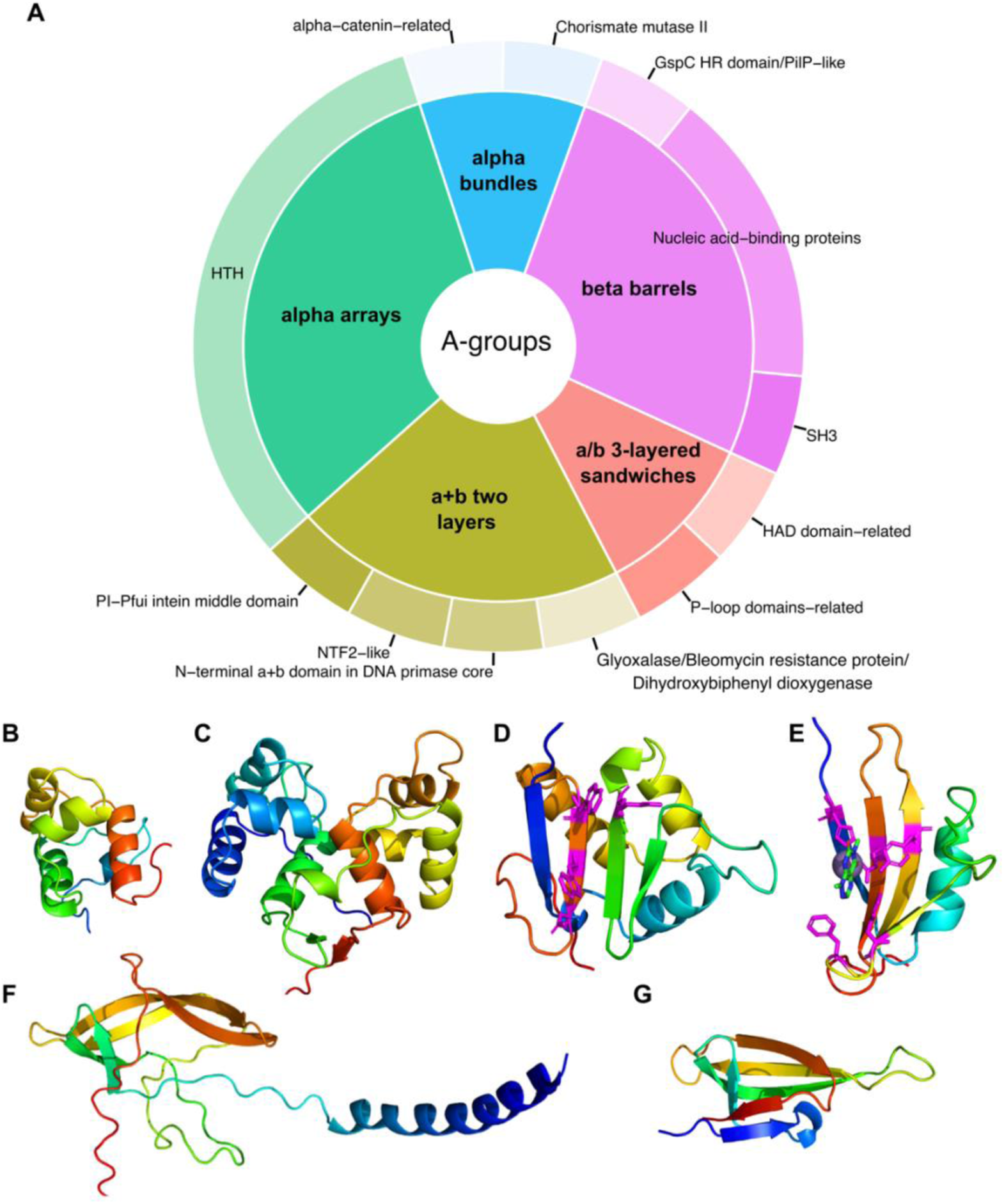
Novel potential pathogenicity island (NPPI7) proteins statistics. **(A)** Distribution of ECOD A- (inner pie chart) and H-groups (outer pie chart) for potentially pathogenic proteins within NPPI7. **(B)** AF model of a helix-turn-helix transcriptional regulator (NCBI: ECO0783223.1). **(C)** AF model of hypothetical protein K5A74_004162 (NCBI: EIB4898420.1). **(D)** AF model of hypothetical protein AAK88_22130 (NCBI: ECO0783211.1). Residues that potentially can coordinate small molecules are shown in sticks and colored magenta**. (E)** Domain of toxoflavin lyase from *Paenibacillus polymyxa* (PDB: 3PKX). Residues that bind toxoflavin are colored in magenta. **(F)** AF model of hypothetical protein QD83_004322 (NCBI: EDV2320831.1). **(G)** Structure of translation initiation factor 5A (PDB: 1BKB).

## Materials and Methods

### Identification of potentially pathogenic proteins in *Salmonella* pangenome

We obtained metadata for 450,571 *Salmonella enterica* genomes from NCBI, most of which lack specific details such as isolation_source, host, host_health_state, and host_disease. 79,758 out of 450,571 genomes have gene annotation and can be confidently categorized based on their metadata into three groups: 14,903 from diseased patients (disease); 55,931 from food, food packages, food process facilities, and other food-related facilities or items (food); 8,924 from the environment such as sewage, pond, and soil (environment). We hypothesized that proteins predominantly found in patient isolates but less so in food and environment might be virulence-associated. Thus, we initially clustered proteins encoded by 79,758 *Salmonella enterica* strains using easy-cluster in mmseq2 with identity 50% and coverage 0.8 and aim to identify protein clusters enriched with proteins from the disease group versus the food and environment ones. Recognizing the presence of redundant isolates could skew the enrichment analysis, we intended to remove proteomes highly similar. The similarity of two proteomes was assessed by tallying the proteins in one proteome that coexisted in the same clusters as proteins from the other proteome and then calculating its ratio by dividing it by the total number of proteins in the proteome. Several cutoffs, namely 0.999, 0.998, 0.995, 0.99, 0.98, 0.97, 0.96, and 0.95, were used to remove redundancy. At each cutoff, after removing the redundant proteomes, we identified clusters enriched with proteins from the disease group against the food and environment groups respectively using a Fisher exact test, with a significance level of Holm-Bonferroni adjusted p < 0.05.

### Domain parsing and classification of the *Salmonella* proteins

We obtained 54,797 clusters from 79,758 proteomes by easy-cluster in mmseq2 with identity 50% and coverage 0.8, of which 11,126 clusters contained proteins from no less than 80 proteomes which is around 0.1% of total proteomes. For obtaining structures of protein clusters, we searched against the AlphaFold Database (AFDB) [43] using the representative sequence of each cluster selected by mmseq2 by default with BLASTP in Diamond 2.0.13 [44]. 9,246 clusters with representative sequences aligned with the AFDB, meeting the criteria of at least 90% identity, above 85% coverage, and a length difference of no more than 31 amino acids. For the 1,880 clusters without matches in the AFDB, we generated their structures using AlphaFold [12], using alignments from HHsearch [45] with parameters -Z 100000 -B 100000. These structures were subsequently segmented and classified into ECOD domains via the domain parser for AlphaFold models (DPAM) with default settings described in [15, 46]. Briefly, DPAM relies on ECOD hits found by HHsuite and Dali [45, 47], as well as inter-residue distances and predicted aligned errors (PAE) of AF models to partition structures into compact domains. Each parsed domain was assigned to the ECOD hierarchy by finding an ECOD parent domain showing the highest DPAM probability for being in the same ECOD topology group (T-group) as the query domain. The DPAM probability was calculated based on sequence and structure similarities evaluated by HHsuite and Dali and the consensus between them. Additionally, domains detected by the DPAM pipeline were further evaluated by the number of secondary structure elements (residues predicted as H, G, or I by DSSP [48] were considered a helix, and residues predicted as E or B by DSSP are considered as a strand) and the completeness of a domain. A domain with less than 3 secondary structure elements and without confident (HHsuite probability and coverage 80%) ECOD hits by HHsuite is considered a "simple topology". A domain that is too short compared to its ECOD parent domain (residues aligned) or the median length of the ECOD T-group (length) it is assigned to is considered a "partial domain". Domains passing these filters are regarded as "low confidence" and cannot be automatically assigned if their DPAM probabilities are below 0.85, while the rest are considered "good domains" with confident assignments.

### Calculation of over and underrepresentation of ECOD H-groups for SPIs and potentially pathogenic proteins

The list of genes that are located within the following *Salmonella* pathogenicity islands (SPIs) was retrieved from the published studies: SPI-1 [24], SPI-2 [24], SPI-3 [49], SPI-4 [50], SPI-5 [51], SPI-6 [30], SPI-7 [52], SPI-8 [53], SPI-9 [54], SPI-10 [55], SPI-11 [56], SPI-12 [57], SPI-13 [58], SPI-14 [58]. Over and underrepresentation of SPIs and potentially pathogenic proteins in ECOD homology groups was calculated as the ratio of observed and expected frequencies. The observed frequency for each ECOD H-group of SPI protein domains was calculated as a ratio of the number of the SPI protein domains assigned to this H-group over the sum of all SPI protein domains assigned to any H-group. The observed frequency for each ECOD H-group of potentially pathogenic protein domains was calculated as a ratio of the number of the potentially pathogenic protein domains assigned to this H-group within the *Salmonella* pangenome over the sum of all potentially pathogenic protein domains assigned to any H-group within the *Salmonella* pangenome. The expected frequency for each ECOD H-group was calculated as the ratio of the total number of domains assigned to this H-group in *Salmonella* pangenome over the total number of domains assigned to any H-group in *Salmonella* pangenome. The significance of the representation was checked using the chi-squared test. P-value < 0.01 was considered significant. Statistical analysis was conducted using the R package, v4.2.1.

### Functional analysis of potentially pathogenic proteins

Gene Ontology (GO) [59] biological processes (BPs) information was retrieved for each protein from UniProt KB [60]. BP GO terms were mapped to GO terms from a Metagenomics slim subset. Overall there are 42 top-level BPs in GO Metagenomics slim subset. One protein can take part in several BPs. Over and underrepresentation of potentially pathogenic proteins in BPs were calculated as the ratio of observed and expected frequencies. The observed frequency in each BP was calculated as a ratio of the total number of the potentially pathogenic proteins in a particular BP over the sum of all potentially pathogenic proteins mapped to any BP. The expected frequency in each BP was calculated as the ratio of the total number of *Salmonella* proteins found for each particular BP to the total amount of proteins mapped to any BP. The significance of overrepresentation was checked using the Chi-square test (P-value < 0.0001 is considered significant).

### Identification of Novel Potential Pathogenicity Islands (NPPIs)

To get the average distance between protein clusters, we calculated the pair-wise distance of proteins in each assembly by counting the minimum number of protein-coding genes between target proteins. The average pair-wise distance of protein clusters was calculated and Multidimensional Scaling (MDS) [61] was used to convert the pair-wise distance matrix to one-dimensional protein localization. Using MDS values for each protein-coding gene from *Salmonella* pangenome we sorted obtained list by MDS values. Genes that are adjacent in this list should also be adjacent in the genome. We used this sorted list to identify clusters of potentially pathogenic proteins that could form novel potential pathogenicity islands (NPPIs) based on the following criteria: a) NPPI should contain no less than 5 potentially pathogenic proteins; b) NPPI should contain no more than 5 nonpathogenic proteins in a row. These criteria enabled us to identify approximately two hundred NPPIs. Verification of NPPIs was conducted using the virulence factor database (VFDB) [35]. For each potentially pathogenic protein, we ran BLAST [22] against the set A of VFDB (4232 proteins). We verified the NPPIs by calculating the over and underrepresentation of potentially pathogenic proteins with significant hits in VFDB within each NPPI, as the ratio of observed to expected frequencies. Any BLAST hit with an E-value < 0.01 was considered significant. The observed frequency for each NPPI was calculated as a ratio of the number of the potentially pathogenic proteins in this NPPI with a significant hit in VFDB over the sum of potentially pathogenic proteins in any NPPI with a significant hit in VFDB. The expected frequency for each NPPI was calculated as the ratio of the total number of potentially pathogenic proteins in this NPPI over the total number of potentially pathogenic proteins in any NPPI. The significance of the representation was checked using the chi-squared test. P-value < 0.01 was considered significant.

## Supporting Information Captions

**Supplementary File 1. List of identified novel potential pathogenicity islands (NPPIs)**

**Supplementary Table S1. Fraction of domains in different ECOD architecture groups**

**Supplementary Table S2. SPIs-related proteins**

**Supplementary Table S3. Novel potential pathogenicity island (NPPI7) proteins**

